# Ageing red deer alter their spatial behaviour and become less social

**DOI:** 10.1101/2021.06.11.448092

**Authors:** Gregory F Albery, Tim H. Clutton-Brock, Alison Morris, Sean Morris, Josephine M Pemberton, Daniel H Nussey, Josh A Firth

## Abstract

Social relationships are important to many aspects of animals’ lives, and an individual’s connections may change over the course of their lifespan. Currently, it is unclear whether social connectedness declines within individuals as they age, and what the underlying mechanisms might be, so the role of age in structuring animal social systems remains unresolved, particularly in non-primates. Here, we describe senescent declines in social connectedness using 43 years of data in a wild, individually monitored population of a long-lived mammal (European red deer, *Cervus elaphus*). Applying a series of spatial and social network analyses, we demonstrate that these declines occur due to within-individual changes in social behaviour, with correlated changes in spatial behaviour (smaller home ranges and movements to lower-density, lower-quality areas). These findings demonstrate that within-individual socio-spatial behavioural changes can lead older animals in fission-fusion societies to become less socially connected, shedding light on the ecological and evolutionary processes structuring wild animal populations.

## Introduction

Identifying the drivers of a wild animal’s social connectedness is important for understanding diverse processes like pathogen transmission^1–3^, information acquisition^4,5^, and fitness^6,7^. In particular, individual ageing provokes broad phenotypic changes, and is therefore likely to impact sociality through a range of mechanisms^8^ (Table 1). Despite growing knowledge concerning senescent changes in wild animals^9,10^, including several examples of senescing behavioural traits^11–13^, relatively little is yet known about the mechanisms underlying age-related declines in sociality (“social senescence”). In recent years, the popularity of social network analysis has facilitated a profound growth in our knowledge of the ecology and evolution of sociality in wild animals^14,15^. Similarly, a recent deeper understanding of senescence in the wild has allowed researchers to identify within-individual declines separate from other demographic processes^9,10^. Combining these two knowledge bases (social networks and senescence in the wild) could lend useful insights into the role of ageing in shaping animal behaviour and structuring wild animal societies, thus informing a wide range of ecological and evolutionary phenomena.

**Table 1.** Potential non-exclusive explanations causing animals to reduce social behaviour as they age. Asterisks denote within-individual changes – i.e., processes that drive “social senescence” *per se* – while mechanisms without asterisks are processes that could drive apparent age-related changes but without necessitating any within-individual changes. NB our data were not able to identify roles of competition avoidance or exclusion.

| Mechanism | Explanation |
| --- | --- |
| **Competition avoidance\*** | Older individuals are less competitive (e.g. due to physiological changes), motivating them to avoid conspecifics. |
| **Exclusion\*** | Older individuals are excluded by younger or more competitive individuals. |
| **Spatial behaviour changes\*** | Older individuals alter spatial behaviours like habitat selection, with their social positions changing as a result. Tested using the SPDE extensions of all model sets, and using the spatial behaviour metrics in model sets 4-6. |
| **Selective disappearance** | More social individuals are more likely to die earlier (e.g. through competition), creating an apparent age-related decline. Tested using model sets 2 and 5. |
| **Demographic changes** | Older individuals' associates die as they age, and are then not replaced. Tested using model set 3. |

Social senescence might occur for many reasons, ranging from individual- to population-level in scale (Table 1). At the individual level, physiological changes may render older individuals less competitive^16^; they may therefore avoid associating with (younger) conspecifics to avoid being outcompeted, or they may be actively excluded. Similarly, ageing individuals may show increased “social selectivity,” replacing disadvantageous or aggressive interactions over the course of their lives with fewer, more positive interactions^12^. In humans, age-related physiological declines (and resulting healthcare requirements) are associated with friends being replaced with family members in the ageing individual’s social network^17^. Alternatively, rather than being driven by physiological decline, such changes could be driven by changing motivations with age – for example, where sociality is beneficial in earlier life but not in old age^18–20^. In species with relatively fluid social systems, an individual’s observed sociality can be influenced by its movement within its environment; in such systems, age-related changes in spatial behaviours could be associated with concurrent changes in observed social connectedness, without actually being driven by a reduction in social behaviour *per se*^14,21^. As such, providing evidence for social senescence may require demonstrating that the observed changes are robust to changes in spatial behaviour. For example, if older individuals have smaller home ranges, they will likely also make fewer unique contacts^21^. Further, ageing individuals may select less desired habitats or prefer areas that other individuals tend to avoid (and which therefore host lower conspecific densities), both of which will likewise reduce social connections^3,14,22^. The relevance of each of these processes will likely depend on the social system of the animal in question: for example, in species with highly stable social groups that move and forage together, it may be unlikely that age-related changes in spatial behaviours underlie changes in sociality.

Notably, observed age-related patterns may not originate from within-individual changes: if certain individuals have higher mortality rates than others, then an apparent age-related pattern might emerge at the population level (i.e., “selective disappearance”)^9,10^. For example, if more social individuals are more likely to die because of greater levels of competition, a population-level pattern of decreasing sociality with age would emerge without requiring any within-individual decline in sociality. Identifying and differentiating selective disappearance from within-individual senescence requires longitudinal analyses following known individuals^9,10^. Similarly, because an animal’s observed sociality depends on surrounding population structure^21,23,24^, demographic changes could produce apparent age-related social declines. For example, if an animal forms relationships when it is young which are not replaced when those contacts die^25,26^, then older animals will be less socially connected as a result. Social connectedness has recently been shown to correlate with age in chimpanzees, macaques, ibex, and marmots^12,27–30^, but the relative roles of these spatial, demographic, and within-individual drivers have yet to be investigated. Untangling these processes could shed a light on the relative importance of within-individual age-related behavioural changes (compared to physiological or demographic processes) in determining population structure in wild animals. Ultimately, doing so will help to inform the role that ageing individuals play in a system’s ecology through processes like pathogen transmission, cooperation, or competition with its associates.

We investigated age-related declines in social behaviour using a wild, long-term study population of individually monitored red deer (*Cervus elaphus*)^31^. The deer, which have been studied since 1973, inhabit a ∼12km^2^ area in the north block of the Isle of Rum, Scotland. They are individually followed from birth, and exhibit a fission-fusion social system characterised by substantial mixing but with repeatable social phenotypes based on grouping patterns^21^. The deer are well-suited to studying social senescence: they experience strong age-related declines in ranging behaviour^11^ and fitness^32^, and they have well-characterised spatially structured social networks structured by a combination of spatiotemporal drivers and individual phenotypes^21^. Importantly, these data are observational, and are not able to differentiate between all the hypothetical drivers of social decline that we outline in Table 1. Nevertheless, by fitting a series of spatially explicit network models, we examine how age-related declines in social behaviour could arise through within-individual senescence, changes in spatial behaviour, and demography, beginning to unpick the potential causes of age-related social changes in wild mammals.

## Results

### Evidencing social declines

Using 43 years of census data containing over 200,000 observations of 935 individually known female deer, we fitted generalised linear mixed models (GLMMs) to investigate the phenotypic drivers of three annual measures of social connectedness: mean group size, degree centrality, and connection strength. We uncovered age-related declines in all three measures when accounting for a range of other intrinsic and extrinsic drivers (**Model Set 1**; Figure 1A-C). Older females had smaller groups, fewer contacts, and weaker social connections (Figure 1A-C). Ageing 1 year came with a reduction in average group size of 0.27 individuals (CI -0.36, -0.21; P<10^−6^; Figure 1A), 0.66 fewer unique contacts (CI -0.83, -0.47; P<10^−6^; Figure 1B), and 0.05 weaker network connection strength (CI -0.07, -0.04; P<10^−6^; Figure 1C).

**Figure 1:**
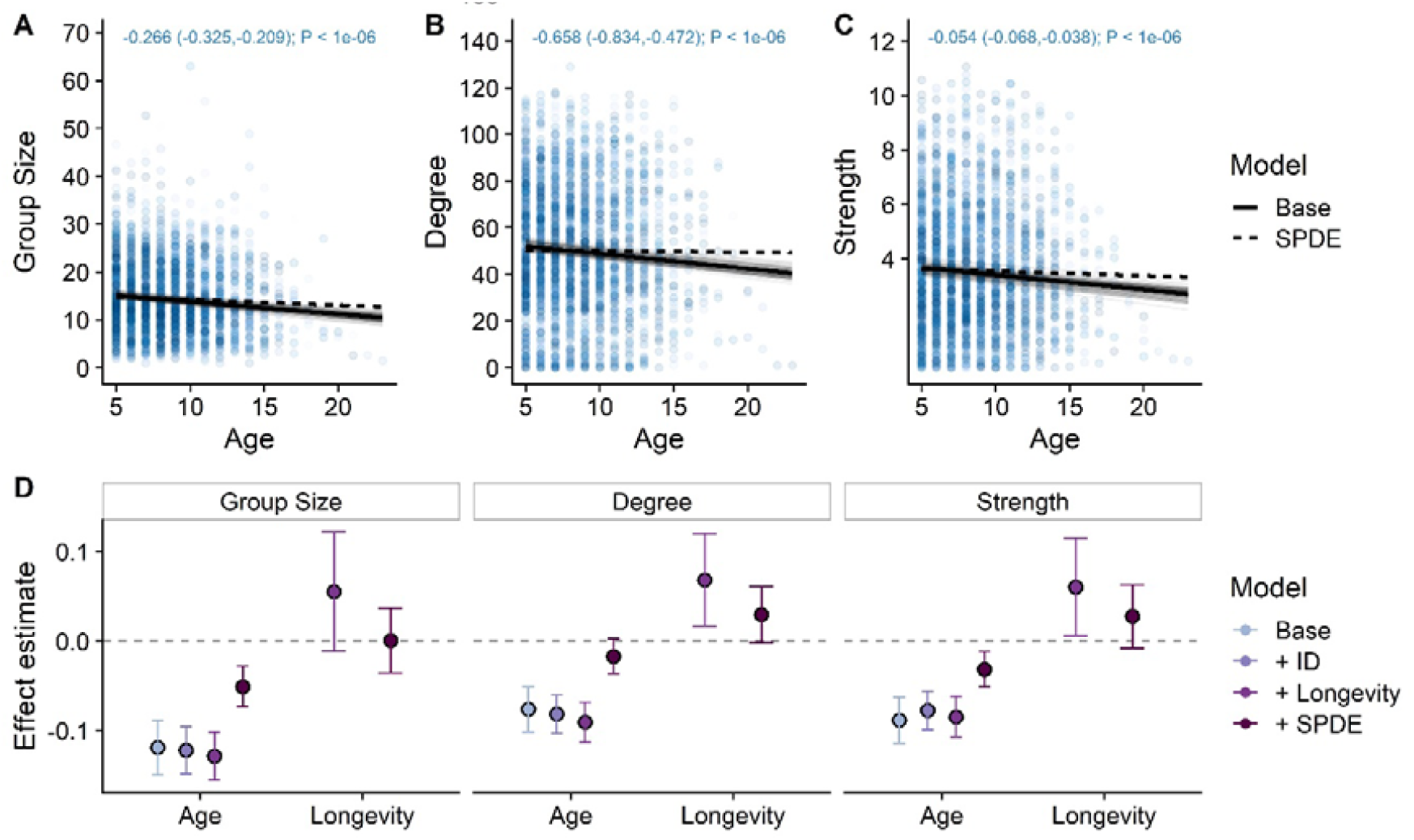
Ageing was associated with reduction in social connectedness. Panels A-C display age-related declines based on 3830 annual observations of 935 individuals, where each point is an observation, with point shading denoting individual age (lighter=older). The text at the top of panels A-C displays the effect estimate (expressed in units per year), with 95% credibility intervals in brackets, and the P value. Opaque black lines are derived from linear model fits from **Model Set 1**, taking the mean of the posterior distribution. Transparent grey lines represent 100 fits drawn randomly from the posterior estimate distributions of each model, to demonstrate error in the slope and intercepts. Dotted black lines display the effect for the SPDE model, demonstrating the result’s robustness to controlling for spatial autocorrelation. Panel D displays the model effect estimates taken from **Model Set 2** (N=3242; effects expressed in units of standard deviations) for age and longevity effects for each social behaviour trait with increasingly complex model formulations, demonstrating that selective disappearance was not responsible for the age-related declines in social connectedness. Dots represent the mean of the posterior effect estimate distribution; error bars denote the 95% credibility intervals of the effect. NB each additional model includes the variables of the models above it (e.g., the longevity model also includes the ID random effect). See Supplementary Table 2 for full effect estimates.

**Figure 2:**
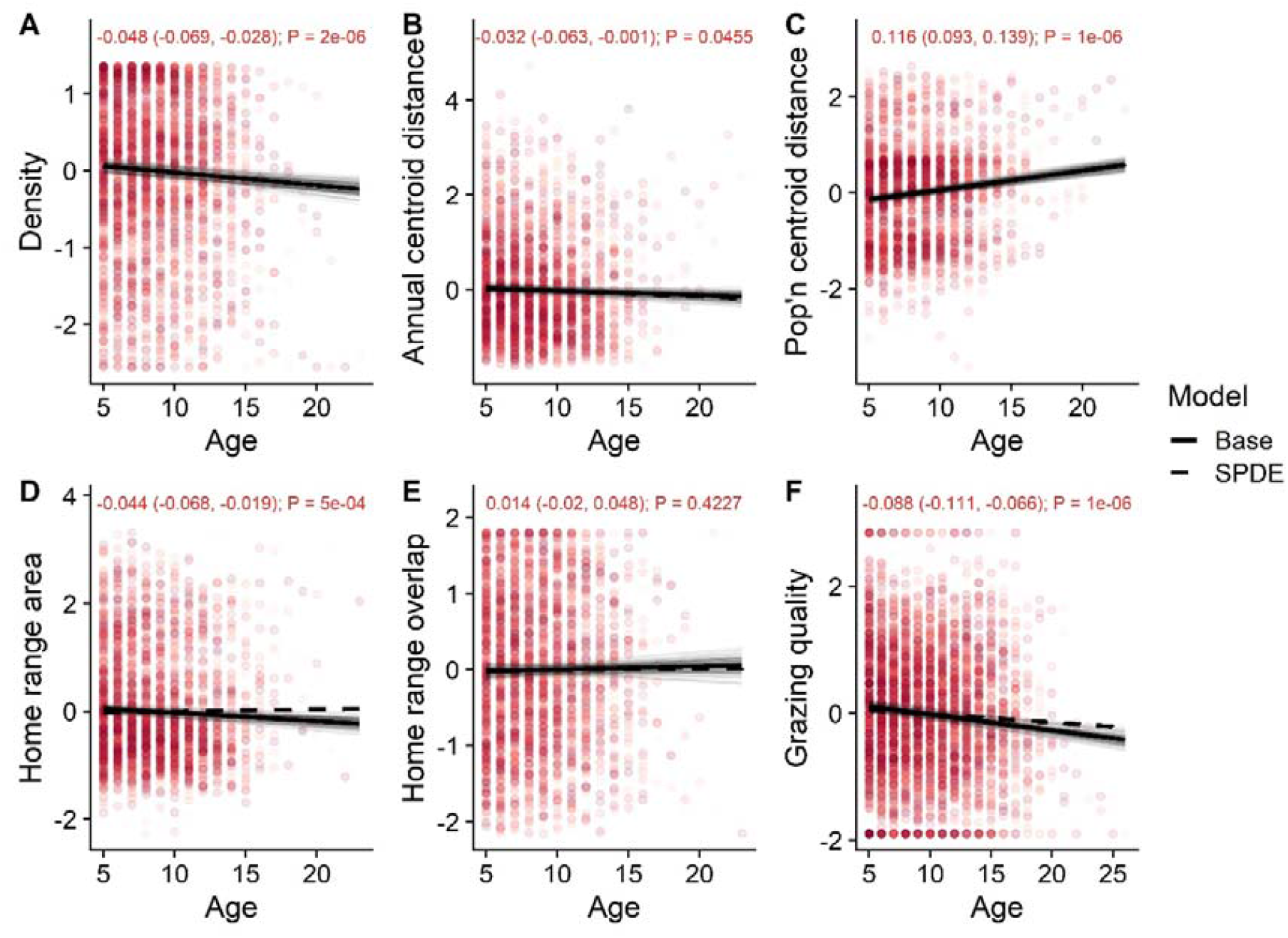
Ageing was associated with changes in a range of socio-spatial behaviours. Each point represents an annual observation of an individual, with point shading denoting individual age (lighter=older). The text at the top of the panels displays the effect estimate for the Base models, with 95% credibility intervals in brackets, and the P value, taken from **Model Set 4**. Opaque black lines are derived from linear model fits, taking the mean of the posterior distribution. Transparent grey lines represent 100 fits drawn randomly from the posterior estimate distributions of each model, to demonstrate error in the slope and intercepts. Dotted black lines display the effect for the SPDE model, demonstrating the result’s robustness to controlling for spatial autocorrelation. The y axes are in units of standard deviations, and were centred around the mean.

### Testing selective disappearance

An age-related decline in social connectedness could be produced if highly social individuals are more likely to die (i.e., “selective disappearance”), creating an apparent population-level decline in sociality. To test this possibility, we sequentially added individual identity and longevity (age at death) to a model constructed on the subset of individuals with known death year (90% of individuals; **Model Set 2**). Longevity was positively associated with degree centrality (0.068; CI 0.017, 0.12; P=0.01) and connection strength (0.06; CI 0.006, 0.115; P=0.03), indicating that individuals with more and stronger contacts were likely to live longer (Figure 1D). However, incorporating individual identity and longevity did not notably change or remove the negative age effect estimates, demonstrating that the observed senescence did not originate from selective disappearance of more-social individuals (P<10^−6^; Figure 1D).

### Testing demographic drivers

We tested whether the death of associates could be driving age-related social declines by examining the relationship between an individual’s summed social connections to individuals that had died in the preceding year, and her own sociality in the focal year (**Model Set 3**). We found that the death of associates was not predictive of social metrics, regardless of whether these associates had died naturally or not (Supplementary Table 1; ΔDIC>0.04). This finding implies that mortality-associated loss of associates is offset by the acquisition of new connections with surviving individuals.

### Examining spatial autocorrelation

Accounting for spatial positioning is important in social network ecology because spatial autocorrelation may produce confounding between response and explanatory variables, potentially rendering analyses less conservative^33^. Specifically, in social network analyses, individuals that live in closer proximity to one another may exhibit more similar social phenotypes because they experience similar environments, and they socialise with each other more frequently^21^. We fitted Stochastic Partial Differentiation Equation (SPDE) effects to consider spatial autocorrelation by accounting for spatial variation in the response variable across the landscape (see Methods). We found that fitting the SPDE effect substantially improved the fit of our models (ΔDIC>353) and reduced age effect estimates in **Model Set 1-2** (Figure 1); the age effects remained significant for group size (P=2×10^−6^) and strength (P=0.009), while degree centrality became nonsignificant (P=0.45). The SPDE effect also removed both significant longevity effects in **Model Set 2** (P>0.07; Figure 1; See Supplementary Table 2 for full effect estimates). Age was heavily spatially structured (ΔDIC=269): individuals with more similar ages lived in closer proximity, with younger individuals in the centre of the study area, and with age increasing to the fringes of the population (Figure 3A). This trend implied that age-dependent changes in spatial behaviour could be contributing to the observed declines in social connectedness.

**Figure 3:**
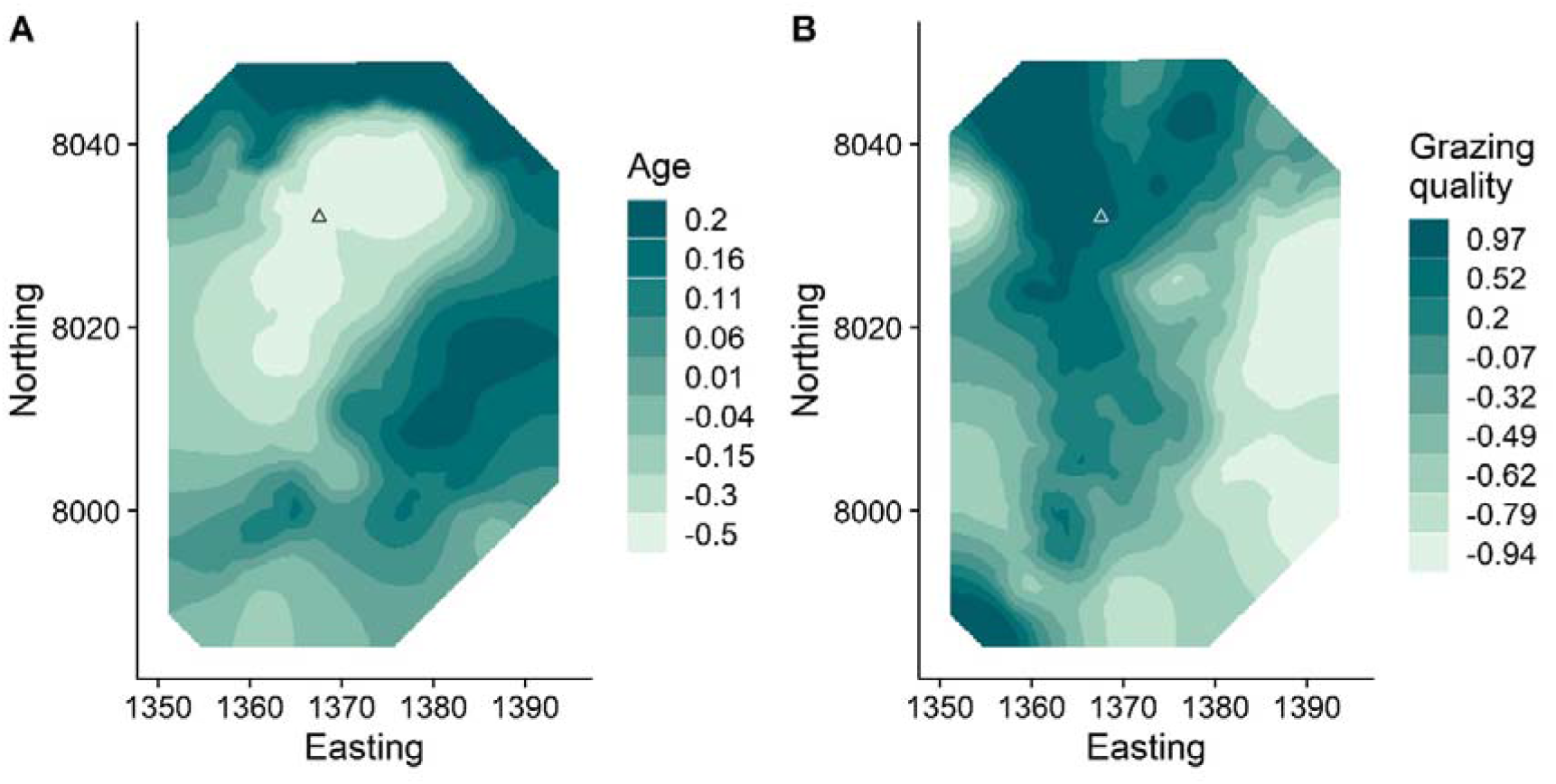
The spatial distributions of age of mature females (A) and grazing quality (B) within the study area. Darker colours relate to greater age or grazing quality. Both figures were obtained by plotting the two-dimensional distribution of the SPDE random effect in INLA GLMMs, with age and grazing quality as response variables respectively. Triangle points represent the population’s centroid, obtained by taking the average location of all individuals’ annual centroids. Ten axis units = 1KM.

### Evidencing declines in spatial behaviour metrics

We quantified age-related changes in a correlated suite of eight annual spatial behaviour metrics, to provide a general assessment of changes in individuals’ spatial behaviour through their lives (**Model Set 4**; Figure 2; See Supplementary Table 3 for full effect estimates). These metrics included: local population density; distance between annual centroids (i.e., mean X and Y coordinates); distance from the population centroid; home range area; proportional overlap with prior annual home range; and average grazing quality. Older deer occurred in lower-density areas (−0.048; CI -0.069, -0.028; P=2×10^−6^; Figure 2A), further from the centre of the study area (0.116; CI 0.093, 0.139; P<10^−6^; Figure 2C), and moved their annual centroids slightly less between years (−0.032; CI -0.063, -0.001; P=0.045; Figure 2B). Older individuals also had smaller home ranges (−0.044; CI -0.068, -0.019; P=5×10^−4^; Figure 2D), but this result did not hold when accounting for spatial autocorrelation in home range size (0.011; CI -0.003, 0.026; P=0.127; Figure 2D). There was no age trend in the degree of overlap between consecutive annual home ranges (0.028; CI -0.005, 0.061; P=0.1; Figure 2E), implying that home ranges do not shrink within themselves. Older individuals were less likely to be observed on high quality grazing (−0.088; CI - 0.111, -0.066; P<10^−6^; Figure 2F).

### Testing selective disappearance for spatial behaviour metrics

Following our protocol for social metrics (**Model Set 2**), we added longevity to investigate how selective disappearance could affect observed spatial metrics’ relationships with age (**Model Set 5**; Supplementary Figure 1). Longer lifespan was associated with those individuals that: inhabited areas of lower density (−0.068; CI -0.102, -0.033; P=0.0002; Supplementary Figure 1A); moved further between annual centroids (0.046; CI 0.007, 0.085; P=0.02; Supplementary Figure 1B); and lived further from the population centre (0.043; CI 0.001, 0.084; P=0.04; Supplementary Figure 1C). Individuals with larger home ranges had shorter lives only when we accounted for spatial autocorrelation (−0.09; CI -0.17, -0.009; P=0.03; Supplementary Figure 1F), and controlling for spatial autocorrelation also removed the longevity effect of annual centroid distance (P=0.87; Supplementary Figure 1B). Neither home range overlap nor grazing quality were associated with longevity (P>0.7; Supplementary Figure 1D-E). Notably, controlling for selective disappearance did not decrease or remove any age effects except that of local density (P=0.12; Supplementary Figure 1A).

### Testing spatial explanations for social declines

We then fitted spatial metrics as explanatory covariates to test whether they could explain age-related declines in social metrics (**Model Set 6**). We found that social metrics were positively associated with local density and home range area, agreeing with previous findings^21^. Additionally, social metrics were negatively associated with distance from the population centre (P Value<10^−6^). Fitting these effects as explanatory variables in the GLMMs did not fully supplant the negative effects of ageing (P<0.041; Supplementary Figure 2; Supplementary Table 4), although it did substantially reduce the size of the estimated effects (Supplementary Figure 3). Including the SPDE effect alongside these three spatial covariates removed the significant associations of age with degree and strength (P>0.15), but retained an association with group size (P Value=0.0002; Supplementary Figure 2B; Supplementary Figure 3).

## Discussion

These observations demonstrate senescent declines in social connectedness in a wild ungulate, while providing much-needed insights into the potential underlying drivers. We uncovered no evidence for selective disappearance or demographic mechanisms governing age-related declines in social connectedness; instead, our results suggest that such declines occurred at the within-individual level. We found that generalised spatial phenotypes also changed with age, as represented by a correlated suite of spatial metrics: older individuals were generally found with smaller home ranges, further from the centre of the population, in areas of lower density, and with lower-quality grazing. This altered spatial behaviour explained age-related declines in degree centrality and connection strength, but not group size, implying the partial involvement of a non-spatial driver of social senescence. Moreover, due to the reciprocal relationships between spatial and social behaviour^14,34,35^ and our use of an observational dataset, it is possible that the changes in spatial behaviour arose from the changes in social connectedness, rather than *vice versa*. Nevertheless, these observations support the value of examining spatial context when considering the intrinsic drivers of social network structure^3,14,21^, particularly when combining long-term data with spatially explicit network models. Social senescence could emerge from generalised reductions in spatial activity caused by physiological decline and inhibited movement ability^11^. This mechanism would be supported if older individuals’ ranges gradually shrank within themselves, while moving less in space. This explanation was not supported for several reasons: first, although an age-related decline in home range area was evident (supporting earlier findings^11^), it was not robust to controlling for spatial autocorrelation, which implies that this finding could originate from age-related changes in spatial locations or spatially structured observation effort rather than smaller home ranges *per se*. Second, older individuals’ home ranges did not increasingly fall within their previous home ranges, implying that they inhabited different areas rather than smaller subsets of the same area. Third, age-related declines in shifts in average annual location were very weak and not robust to controlling for spatial autocorrelation, demonstrating that home ranges shifted at a relatively constant rate across the landscape through an individual’s life rather than notably slowing down in old age. Taken together, these observations imply that ageing females exhibit similar spatial activity levels but in different areas of the landscape.

Although reduced activity did not appear to be responsible, altered spatial behaviour did contribute to age-related declines in social connectedness. Ageing individuals were less social partly because they preferred to inhabit lower-density, lower-quality areas at the edge of the study area which offer fewer social opportunities. They may ultimately inhabit these areas because physiological changes cause ageing individuals to alter their habitat selection. Red deer teeth are worn down as they age^36,37^, reducing their ability to ingest food as efficiently. Older deer may accommodate these physiological changes by moving to areas that allow them to feed on alternative vegetation (e.g. if longer grasses are easier to crop with worn incisors), which also contain lower densities of conspecifics. In this way, social declines could arise partly as a by-product of habitat selection based on physiological ageing rather than being related to changes in social preferences itself.

Nevertheless, we found that social senescence was evident even when spatial factors were accounted for: older individuals still had smaller groups even when considering their home range areas, local population density, and location on the landscape, implying that alterations in spatial behaviour could not be the only explanation for the deer becoming less social with age. Although we tested six spatial metrics and three social metrics, these behaviours are extremely unlikely to be independent: for example, moving away from the centre of the population, towards areas of lower density, and having fewer social associates are all likely to be part of the same correlated selection of behaviours that are very difficult to discern using observational data. Furthermore, while we fitted spatial behaviours as response variables in our models of social connectedness, social and spatial behaviour are part of a bidirectional process involving a number of feedbacks^34,35,38^. As such, although we formulate spatial behaviours as being explanatory of changes in social connectedness, it is possible that changes in spatial behaviour were themselves influenced by changes in social behaviour. Our results should not necessarily be interpreted as separate evidence for (absence of) changes in each spatial behaviour in isolation, but to indicate that age is associated with generalisable within-individual changes in socio-spatial behavioural syndromes and reduced social connectedness.

Social senescence could be a response to aggressive or competitive interactions with younger individuals that may have greater resource demands: to give an example, reproductive female olive baboons (*Papio anubis*) are more aggressive to other females for this reason^39^. Older female deer are less likely to reproduce^32^, but reproductive status was included in our models and is therefore unlikely to be directly responsible for the observed social senescence. Alternatively, ageing deer may reduce their connections to individuals with whom they have had more aggressive interactions in favour of more positive interactions, thereby becoming more “socially selective” as they age^12^. This tendency to avoid aggressive interactions could also cause them to have reduced grazing quality (as we observed) if younger individuals monopolise the higher quality resources, or if the younger individuals are more competitive and older individuals avoid this competition by moving to less-favoured areas (i.e., competitive exclusion). This explanation was supported by a longevity effect in high-density areas, which is congruent with competition-related mortality driven by insufficient local resources. Inferring such competitive or aggressive interactions would likely require high-resolution methods like direct behavioural observation in combination with telemetry, which provide detailed information on fine-scale movement patterns but which often currently come with important restrictive tradeoffs in terms of how many individuals are GPS-tagged or observed and for how long^40,41^. Telemetry- or behavioural observation-based approaches may not be feasible to run for the duration necessary to detect the subtle, life-long patterns of social senescence we observed. Crucially, our long-term longitudinal data (complete with high-certainty life history measures) were also able to differentiate selective disappearance and changes in demographic network structure from within-individual declines, which may be important for producing age-related social changes in other populations (e.g. ibex^29^). These findings therefore imply a tradeoff between the resolution and breadth of behavioural data, while supporting the value of individual-based long-term studies of wild animals for identifying senescence^9^ and behavioural changes^29^.

Older individuals may decrease their social connectedness due to changing motivations as they age – for example, if it is advantageous to be well-connected when young (e.g. to gain social information) but less so when older (e.g. due to previously accumulated knowledge)^18–20^. Alternatively, if older individuals undergo immunosenescence^42^ they may reduce social connections as a compensatory measure to avoid parasite exposure. Identifying these kinds of mechanisms would require investigating the fitness- and disease-related consequences of sociality, asking whether the fitness of younger individuals holds a different relationship to social connectedness than the fitness of older individuals does. Importantly, such sociality-fitness relationships are likely to depend on a species’ social system; because the deer exhibit a loose, fission-fusion system, our results may be less applicable to species with tight, consistent social groups that move and forage together. Investigating demographic structuring, selective disappearance, and spatial behaviours in primate populations (e.g.^12,27,43^) may shed an important light on the different drivers of age-related changes in sociality across mammals and other taxa. Ultimately, broadening the selection of study organisms for age-sociality interactions to include those a wider variety of fluid and rigid social systems could help to develop generalisable insights about the causes and consequences of social ageing across the tree of life.

In sum, we provide evidence of social senescence in wild deer, and show that this senescence could be explained partly by altered spatial behaviour. Although experiments might be necessary to fully discern the specific drivers of social senescence, we were able to use long-term observational data to successfully identify a within-individual decline (i.e., senescence) in social connectedness that was robust to selective disappearance, demographic changes, and changes in spatial behaviours. As such, these findings provide novel insights into the ecology and evolution of social behavioural changes and the ecological role played by ageing individuals, forming a foundation for understanding how age governs the structure of wild animal societies by shaping individual behaviour.

## Methods

### Data collection and study system

The study was carried out on an unpredated long-term study population of red deer on the Isle of Rum, Scotland (57°N,6°20′W). The natural history of this matrilineal mammalian system has been studied extensively^31^, and we focussed on females aged 5+ years, as these females have the most complete associated census data, few males live year round in the study area, and nearly all females have bred for the first time at the age of 5. These individuals are known to exhibit age-related declines in a range of behavioural and life history traits^11,32,44^. Individuals are monitored from birth, providing substantial life history and behavioural data, and >90% of calves are caught and tagged, with tissue samples taken^31^. Census data were collected for the years 1974-2019, totalling 201,848 census observations. Deer were censused by field workers five times a month, for eight months of the year, along one of two alternating routes^31^. Individuals’ identities, locations (to the nearest 100M), and group membership were recorded. Grouping events were estimated by experienced field workers according to a variant of the “chain rule” (e.g.^45^), where individuals grazing in a contiguous group within close proximity of each other (under ∼10 metres) were deemed to be associating, with mean 204.5 groups observed per individual across their lifetime (range 1-853). If they breed female deer on Rum give birth to a single offspring, typically in May, and we therefore consider their annual cycle as running from May in one calendar year to May the next. All individuals are deemed to be age “zero” in their first year, and all turn “one” on the first of May the following year, regardless of when they were born. This age assignment continues throughout their lives (e.g., in the 2015 deer year, all individuals that were born in 1994 were considered “21 years old”). Accordingly, we summarised individuals’ behaviours based on a “deer year”, which runs from 1^st^ May to 30^th^ April; any female that died in the course of this period was removed. Our full dataset included 3830 observations of 935 individuals. Previous work shows that gestation and lactation impose costs to subsequent survival and reproduction^46,47^. To characterise each female’s investment in reproduction the previous year, we used three categories based on field observations: None (did not give birth); Summer (the female’s calf died in the summer, before 1^st^ October); and Winter (the female’s calf survived past 1^st^ October).

### Deriving behavioural metrics

All code is available at https://github.com/gfalbery/Lonely-Old-Deers. Following previous methodology^21^, we constructed a series of 43 annual social networks using “gambit of the group,” where individuals in the same grouping event (as described above) were taken to be associating^48^. Dyadic associations were calculated using the “simple ratio index”^49^ derived as a proportion of total sightings (grouping events) in which the focal individuals were seen together: Sightings_A,B_/(Sightings_A_+Sightings_B_-Sightings_A,B_). In this dyadic matrix, 0=never seen together and 1=never seen apart. Using the annual social networks, we derived three individual-level annual network metrics that are commonly used across animal social networks and have been considered in detail^24,25,50,51^.

Our measures included three “direct” sociality metrics, which only consider an individual’s connections with other individuals:

1. Group Size – the average number of individuals a deer associated with per sighting;
2. Degree – the number of unique individuals she was observed with over the course of a year, irrespective of how frequently she was observed with them;
3. Strength – the sum of all their weighted social associations to others, also known as “weighted degree”.

Each metric was fitted as a response variable in separate Model Sets. These metrics were well correlated (Supplementary Figure 4), and therefore should not be considered substantially different phenotypes; instead, we present all three metrics under the expectation that they would all change in similar directions, being highly correlated.

We derived 6 correlated metrics related to population structure and spatial behaviour, hereafter referred to collectively as “spatial metrics”. These metrics were to be used first as response variables to identify the drivers of spatial behaviours (**Model Sets 4-5**), and as explanatory variables to identify how spatial behaviours affect social metrics (**Model Set 6**).

1. Local population density (hereafter “**density**”) was calculated based on an annual population-level space use kernel derived in AdeHabitatHR^52^ and using each individual’s annual centroids (i.e., mean X and Y coordinates), following previous methodology^21^. Briefly, for each year, we took each individual’s annual centroid and calculated the spatial density of these centroids (i.e., individuals per KM^2^). Each individual was then assigned a local density value based on their location on this annual kernel.
2. Distance between an individual’s annual centroids was taken to indicate shifting in average location from year to year, to identify whether individuals’ locations move more in their later years (“**Annual centroid distance**”).
3. To detect long-term patterns of an individual’s space use on the two-dimensional landscape, and to identify whether ageing individuals move gradually outwards, we took distance from the overall centre of the population, calculated by taking the mean of all annual centroids’ easting and northing (“**Population centroid distance**”).
4. To examine changes in range size over the course of an individual’s lifespan, we used the 70% isocline of an individual’s space use kernel (“**Home range area**”). Based on each individual’s annual sighting locations and using the AdeHabitatHR package^52^, this was calculated using a density kernel estimator based on previous methodology^11,21^.
5. To investigate the degree to which an individual’s annual home range fell within its previous home range, we used an asymmetrical measure of home range overlap from year to year, using the space use kernels (“**Home range overlap**”). A value of 1 for this variable would indicate that an individual’s home range in year t fell entirely within its home range in year t-1; a value of 0.5 indicates that half of its home range lay within its previous home range; and a value of 0 indicates no overlap with its previous annual home range. Combined with the “Annual centroid distance” metric, this home range overlap metric aimed to test whether an individual’s range generally shrunk to smaller areas within its known range, consistent with physiological decline, rather than an active movement to areas outside its previous range.
6. To investigate changes in grazing quality, for each individual we calculated the proportion of sightings that occurred on high-quality short or long greens each year (“**Grazing quality**”). A value of 1 denoted that an individual was only ever present on these high-quality grasses, while a value of 0 denoted an individual was never seen grazing on them, and was instead always seen on other land types like rock, sand, or lower-quality grazing like heather or *Molinia*.

For metrics that represented year-to-year changes (“Annual centroid distance” and “Home range overlap”), we only included observations that occurred in sequential years (year t and year t+1, year t+1 and year t+2, etc). For example, if a female was only observed in two non-sequential years (year t and year t+2), both timepoints were coded as missing. Importantly, because these variables were well correlated (Supplementary Figure 4), we choose not to interpret these results independently, but to paint a general picture of age-related changes in spatial behaviours, with the caveat that the results of these models are likely non-independent.

### Statistical analysis

To investigate phenotypic associations with social behaviour, we fitted Generalised Linear Mixed Models (GLMMs) using the Integrated Nested Laplace Approximation (INLA) in the R package R-INLA^53,54^. This method allows fitting of a Stochastic Partial Differentiation Equation (SPDE) random effect to control for and quantify spatial autocorrelation^55,56^. The SPDE effect models spatial autocorrelation by estimating to what degree points that are closer to each other in space are more similar than points that are further away^33^. The model then accounts for this similarity, finding sources of variation that occur over and above that expected given the spatial patterns of the response variable. It does so using Matern covariance, approximating the continuous Gaussian field of the response variable using a triangulated mesh of connected discrete locations^53^. The fits of equivalent models with and without the SPDE effect were compared using the Deviance Information Criterion (DIC) to investigate the importance of spatial autocorrelation. All P values are derived from estimating the proportion of the marginal distribution for a given effect that overlapped with zero, and multiplying it by 2. This can be thought of as being the probability of drawing a value greater than or lower than zero (depending on the direction of the effect), per a two-tailed test.

All GLMMs used a Gaussian family specification. For all models, we used noninformative priors for the SPDE effects; we assessed covariance of our explanatory variables and their variance inflation factor (VIF) to ensure models fit well and without being overwhelmed by correlations among our predictors (all VIF values < 2). We checked model fit by assessing correlations between predicted and observed values and by ensuring even distributions of the model residuals.

We constructed 6 sets of models designed to test different mechanisms driving age-related changes in sociality (see Model Set details below). All models included the following “base” fixed effects: Year (continuous); annual Population Size (continuous, log-transformed); Number of observations per individual (continuous); Reproductive Status (three categories: No Calf; Calf Summer Death; Calf Survived to October 1st); Age in years (continuous). All continuous explanatory and response variables were scaled (mean=0; standard deviation=1) to help model fitting. All models included random effects of individual ID and observation year.

Before beginning the behavioural analysis, to investigate the spatial autocorrelation of age and to examine its spatial distribution visually, we fitted a model with age as a response variable and with only the SPDE effect as an explanatory variable.

#### Model Set 1: Social metrics and age

Our first Model Set examined whether our sociality metrics changed over individuals’ lifespans, using mean annual measures (N=3581 female years), and including only the base covariates outlined above. We then added a spatial autocorrelation term using an SPDE random effect in INLA, to investigate whether the models were robust to spatial structuring. Comparing the effect of the base model with the spatial model would indicate whether the response variable is spatially structured, and comparing the effect estimates would reveal whether this changed our conclusions. We used the Deviance Information Criterion (DIC) as a measure of model fit; we chose a change in DIC (ΔDIC) of 10 to identify whether the spatial autocorrelation effect improved the model.

#### Model Set 2: Social metrics and selective disappearance

To investigate whether selective disappearance of individuals may influence estimates of age-related social changes (e.g. if more social individuals were selectively lost), we sequentially added a selection of variables into the models. Because these models only used individuals with a known death year, they used a slightly smaller dataset (N=3242 female years). We first fitted a base model without individual identity as a random effect, and we then sequentially added individual identity as a random effect, and then longevity as a fixed effect (i.e., age at death). An explanation of the test for selective disappearance can be found in Van de Pol and Verhulst^10^. Briefly, fitting both longevity and age in the model isolates the effect of within-individual ageing from that of between-individual selective disappearance. If fitting longevity as a covariate altered the size of the age effects, particularly by rendering them insignificant, it would imply important selective disappearance effects (i.e., between-individual differences) rather than within-individual changes^10,11^. Finally, we included the SPDE effect to examine whether longevity effects were robust to spatial structuring.

#### Model Set 3: Investigating demographic associations

Because the social phenotype of any individual is dependent upon not just itself but also on the other individuals within the population, demographic processes, particularly the loss of an individual’s social associates over time, may passively contribute to age-related changes in remaining individuals’ sociality (e.g. through reducing the potential for encounters with previous associates). To investigate whether these loss processes could drive age-related declines, we tested whether social network positions were predicted by the death of the female’s connections the previous year. We fitted this value as an explanatory variable in the models from Model Set 1, using either 1) connections to individuals that were shot, or 2) those that died for any reason. A negative estimate for this effect would imply that age-related declines could be explained by the death of a deer’s previous connections, rather than solely individual-level change.

#### Model Set 4: Spatial metrics

We investigated how age altered spatial behaviour, using our six spatial metrics as response variables (described above). These models included the same covariates as the social behaviour models (Model Set 1). For annual movement and the home range area and overlap metrics, we also fitted the SPDE effect; for density and population centroid distance, the response variables were explicitly spatially distributed on the landscape, so fitting the SPDE effect would be misleading.

#### Model Set 5: Spatial metrics and selective disappearance

We then repeated the selective disappearance protocol used for social metrics (Model Set 2) on our spatial metrics, to investigate whether any estimated age-related changes in spatial behaviour were altered by selective disappearance of certain individuals.

#### Model Set 6: Investigating how spatial behaviour affects the ageing of sociality

Finally, we investigated whether social metrics were driven by variation in spatial behaviour, and whether age-related declines were robust to these spatial drivers. Because several of our spatial measures were well-correlated, we used a model addition procedure to identify which spatial measures best explained social network metrics. We used the Deviance Information Criterion (DIC) as a measure of model fit. Beginning with the base model, we sequentially added each spatial metric and then compared the DIC of these models. The best-fitting variable (i.e., the one that reduced DIC by the most) was kept, and then the process was repeated, until all variables were fitted or no remaining variables improved the model. We used a change in DIC of 10 to identify variables that improved model fit.

## Acknowledgements

We thank NatureScot and its predecessors for permission to work on the Isle of Rum NNR. The field project has been supported by grants mainly from the UK NERC with some additional funding from BBSRC, the Royal Society and ERC. We thank all who have contributed to the maintenance of the project over time, especially Loeske Kruuk. We thank multiple dedicated field workers who have contributed to field data collection, especially Fiona Guinness who collected the first 20 years of census data. GFA was funded by NSF grant number 1414296, and by a Bruce McEwen Career Development Fellowship the Animal Models for the Social Dimensions of Health and Aging Research Network (NIH/NIH R24 AG065172). JAF was supported by BBSRC (BB/S009752/1) and funding from NERC (NE/S010335/1).

## Supplementary Figures

**Supplementary Table 1:** DIC changes associated with including a variable representing “connections to individuals that died the previous year” in the models of social behaviours (**Model Set 3**). “Dead”=strength of connections to individuals that died. “Shot”=strength of connections to individuals that were shot (a subset of “dead”). To be kept in the model, these metrics would have to reduce DIC by a threshold value of -10.

| Response | Variable | ΔDIC | Kept |
| --- | --- | --- | --- |
| Group Size | Dead | 1.72 | 0 |
|  | Shot | 8.81 | 0 |
| Degree | Dead | 6.84 | 0 |
|  | Shot | 16.78 | 0 |
| Strength | Dead | -0.042 | 0 |
|  | Shot | 10.36 | 0 |

**Supplementary Table 2:**
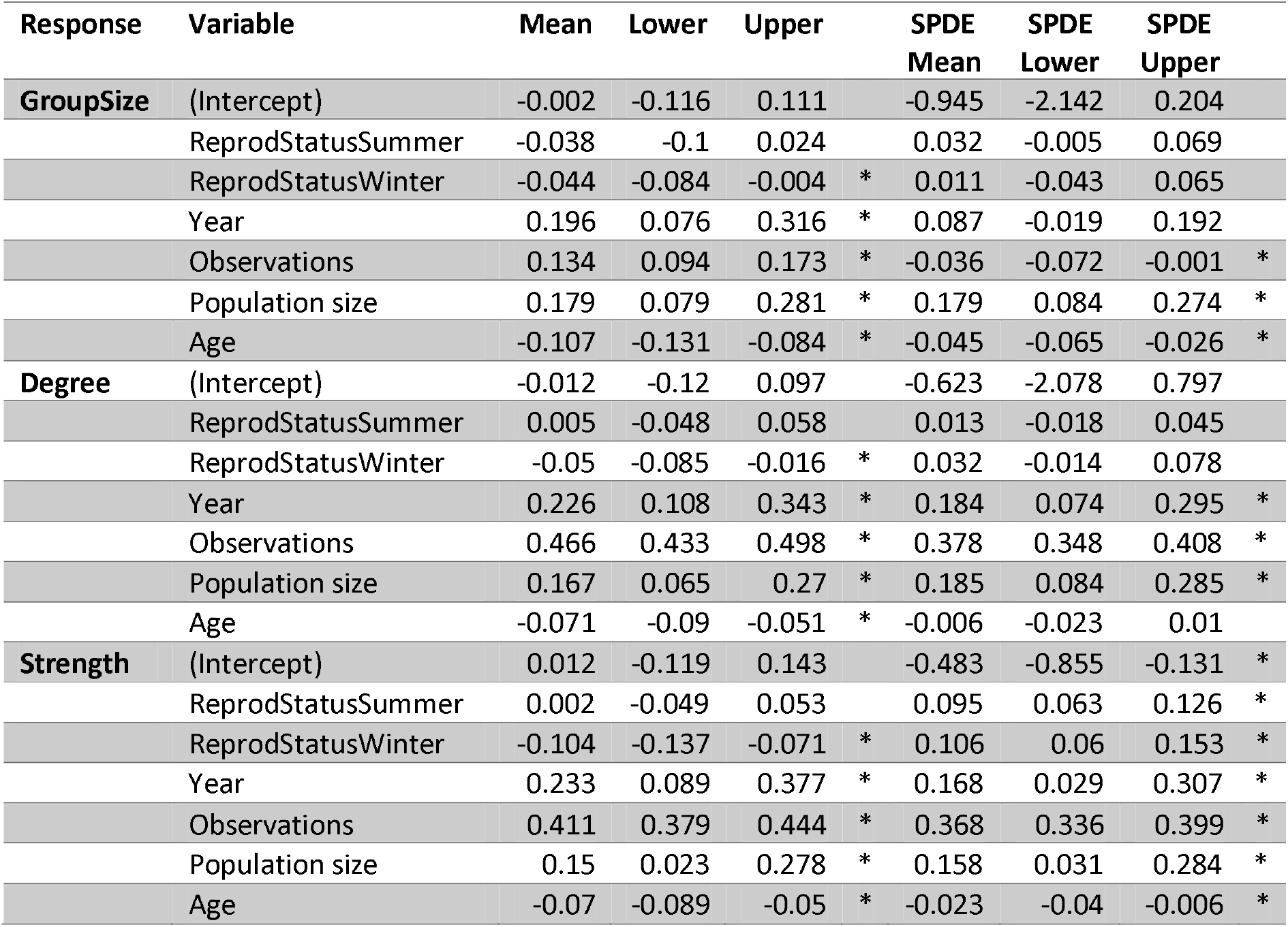
Effect estimates associated with fixed effects in our models of social behaviour (**Model Set 1**). Estimates include the posterior mean and lower and upper 95% credibility intervals, both for the base model and the SPDE model. Asterisks represent significant estimates (i.e., estimates that did not overlap with zero). Estimates are displayed in units of standard deviations, differing from the yearly numbers for raw data changes displayed in Figure 1.

**Supplementary Table 3:**
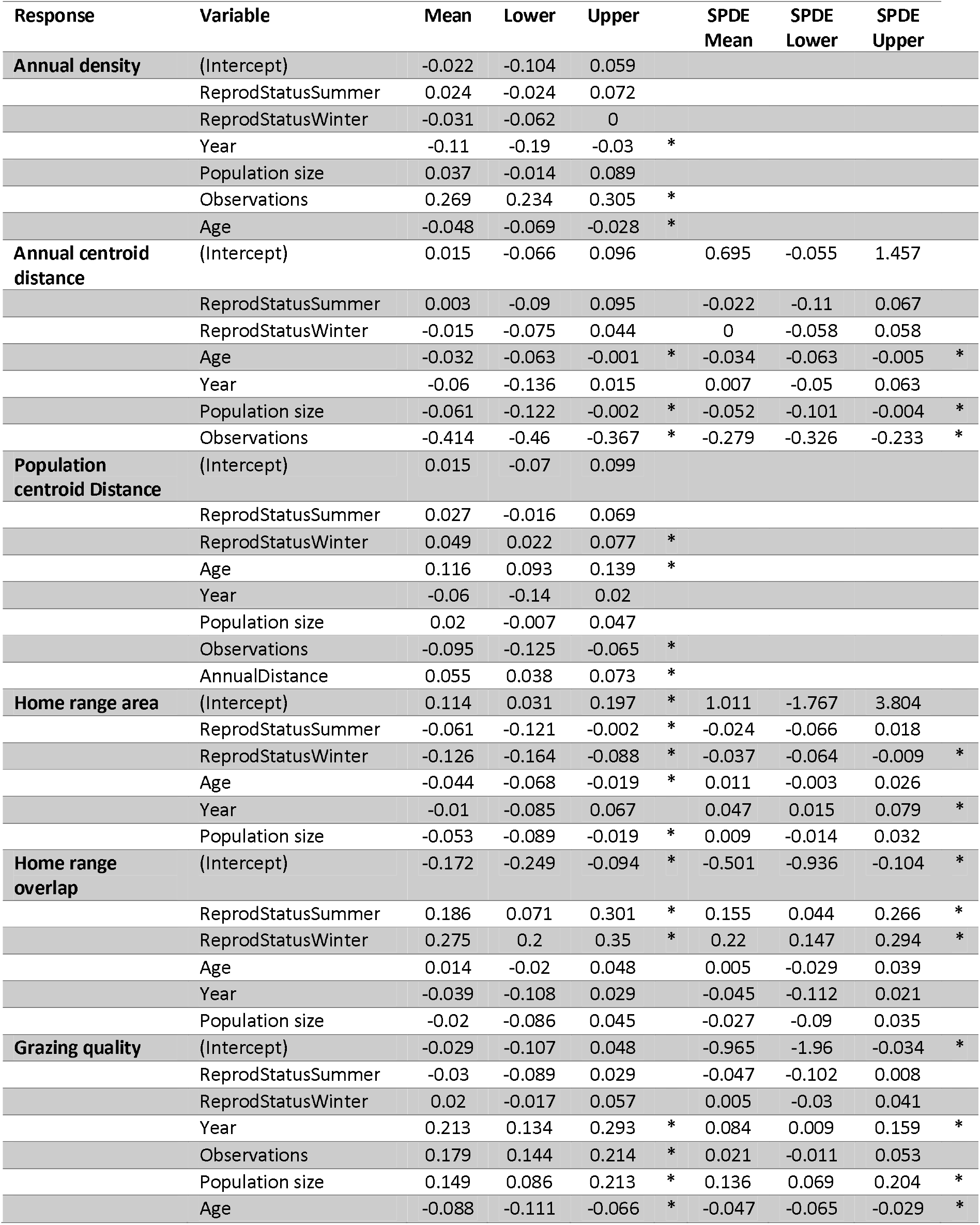
Effect estimates associated with fixed effects in our models of spatial behaviour (**Model Set 4**). Estimates include the posterior mean of the posterior and lower and upper 95% credibility intervals, both for the base model and the SPDE model. Asterisks represent significant estimates (i.e., estimates that did not overlap with zero). Estimates are displayed in units of standard deviations. NB there are no estimates for SPDE effects for annual density and population centroid distance because these are spatially distributed variables for which an SPDE effect was not fitted.

**Supplementary Table 4:** DIC changes associated with adding spatial behaviours as explanatory variables in the models of social behaviours (**Model Set 6**). Variables with asterisks were retained in the round. HRA = Home Range Area.

| Response | Round | Variable | ΔDIC | Kept |
| --- | --- | --- | --- | --- |
| Group Size | 1 | **PopnDistance** | -165.104 | 1 |
|  | 1 | AnnualDensity | -140.133 | 0 |
|  | 1 | HRA | 1.813998 | 0 |
|  | 2 | **AnnualDensity** | -93.2626 | 1 |
|  | 2 | HRA | 1.094292 | 0 |
|  | 3 | **HRA** | -2.18501 | 1 |
| Degree | 1 | **PopnDistance** | -285.413 | 1 |
|  | 1 | AnnualDensity | -182.979 | 0 |
|  | 1 | HRA | -137.134 | 0 |
|  | 2 | AnnualDensity | -79.7089 | 0 |
|  | 2 | **HRA** | -88.2182 | 1 |
|  | 3 | **AnnualDensity** | -211.199 | 1 |
| Strength | 1 | PopnDistance | -109.92 | 0 |
|  | 1 | **AnnualDensity** | -234.801 | 1 |
|  | 1 | HRA | 4.165836 | 0 |
|  | 2 | **PopnDistance** | -49.6076 | 1 |
|  | 2 | HRA | -13.5552 | 0 |
|  | 3 | **HRA** | -2.74231 | 1 |

**Supplementary Figure 1:**
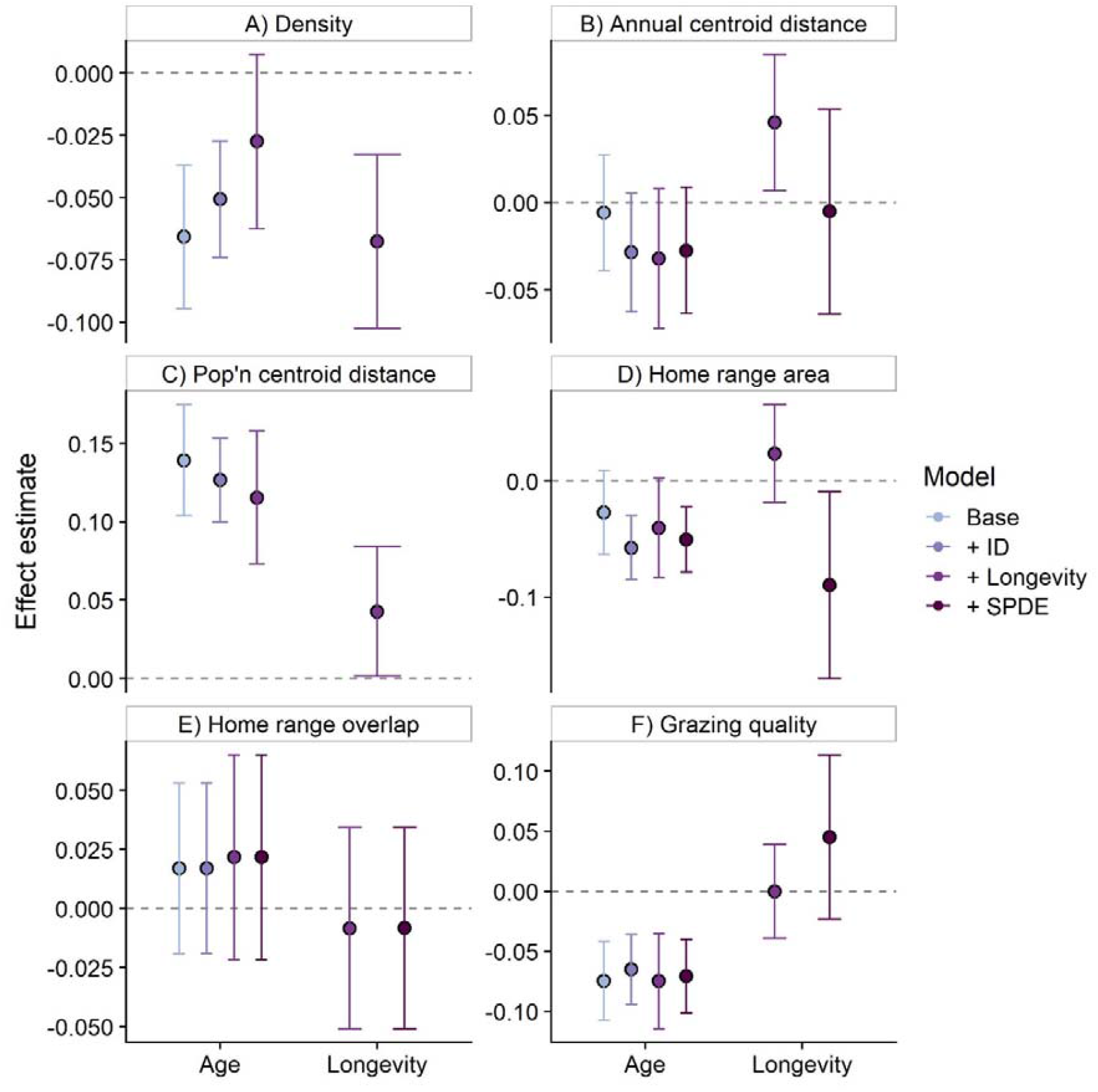
Model effect estimates investigating how selective disappearance drives age-related changes in spatial behaviour (**Model Set 5**). Each panel displays the model effect estimates for age and longevity for each spatial metric, demonstrating that selective disappearance was not responsible for the age-related changes in spatial behaviour, except in the case of density (A). Dots represent the mean of the posterior effect estimate distribution; error bars denote the 95% credibility intervals of the effect. The y axis is in units of standard deviations. NB density and centroid distance did not have an SPDE effect fitted to control for spatial autocorrelation because they are deterministically distributed in space, such that the SPDE effect would be uninformative.

**Supplementary Figure 2:**
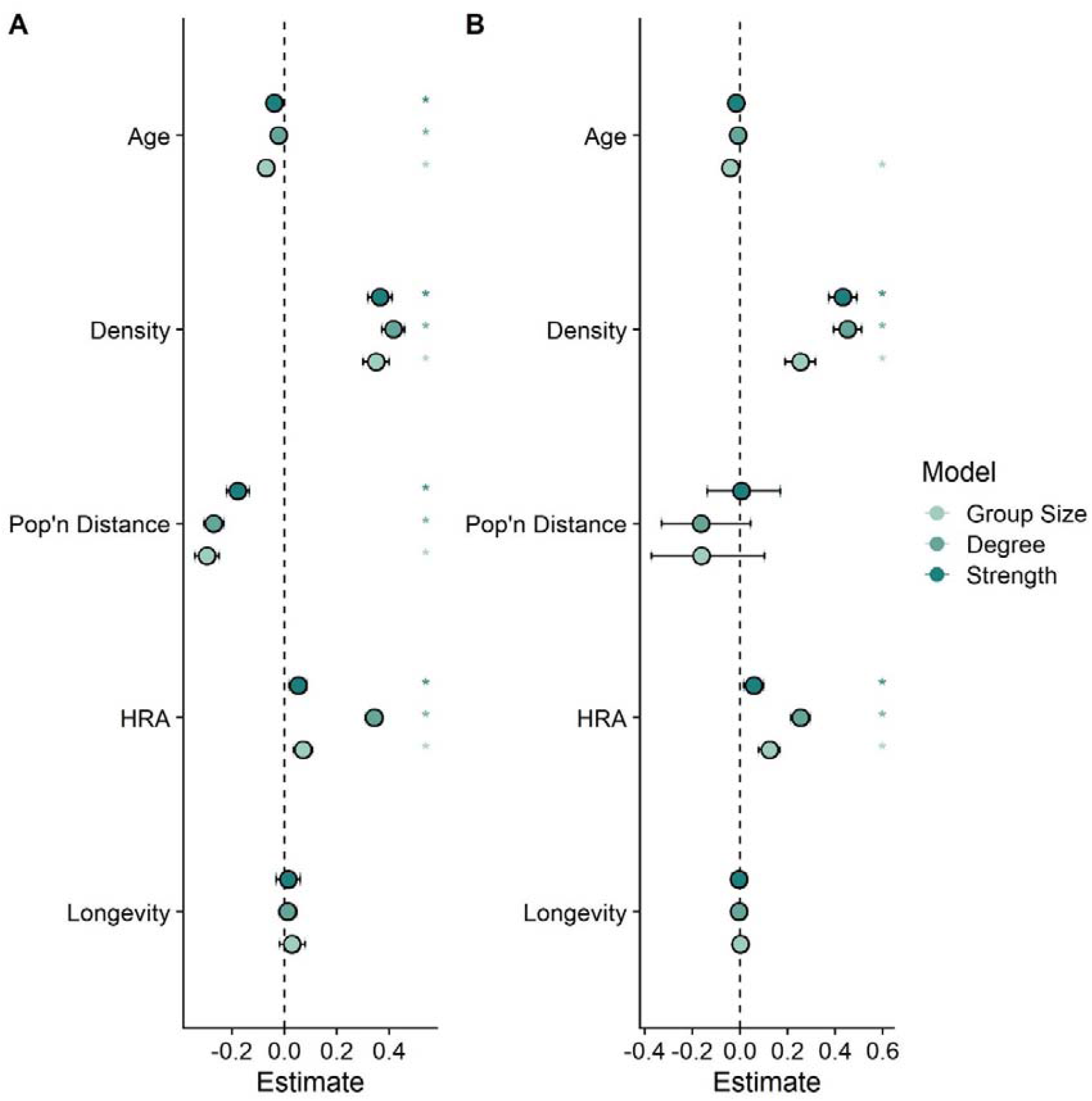
Model estimates describing associations between spatial behaviours and social behaviours, and age-related declines in social behaviour when these variables were accounted for (**Model Set 6**). Dots represent the mean of the posterior effect estimate distribution; error bars denote the 95% credibility intervals of the effect. These estimates are displayed both for the base model (A) and the SPDE model (B). HRA = home range area.

**Supplementary Figure 3:**
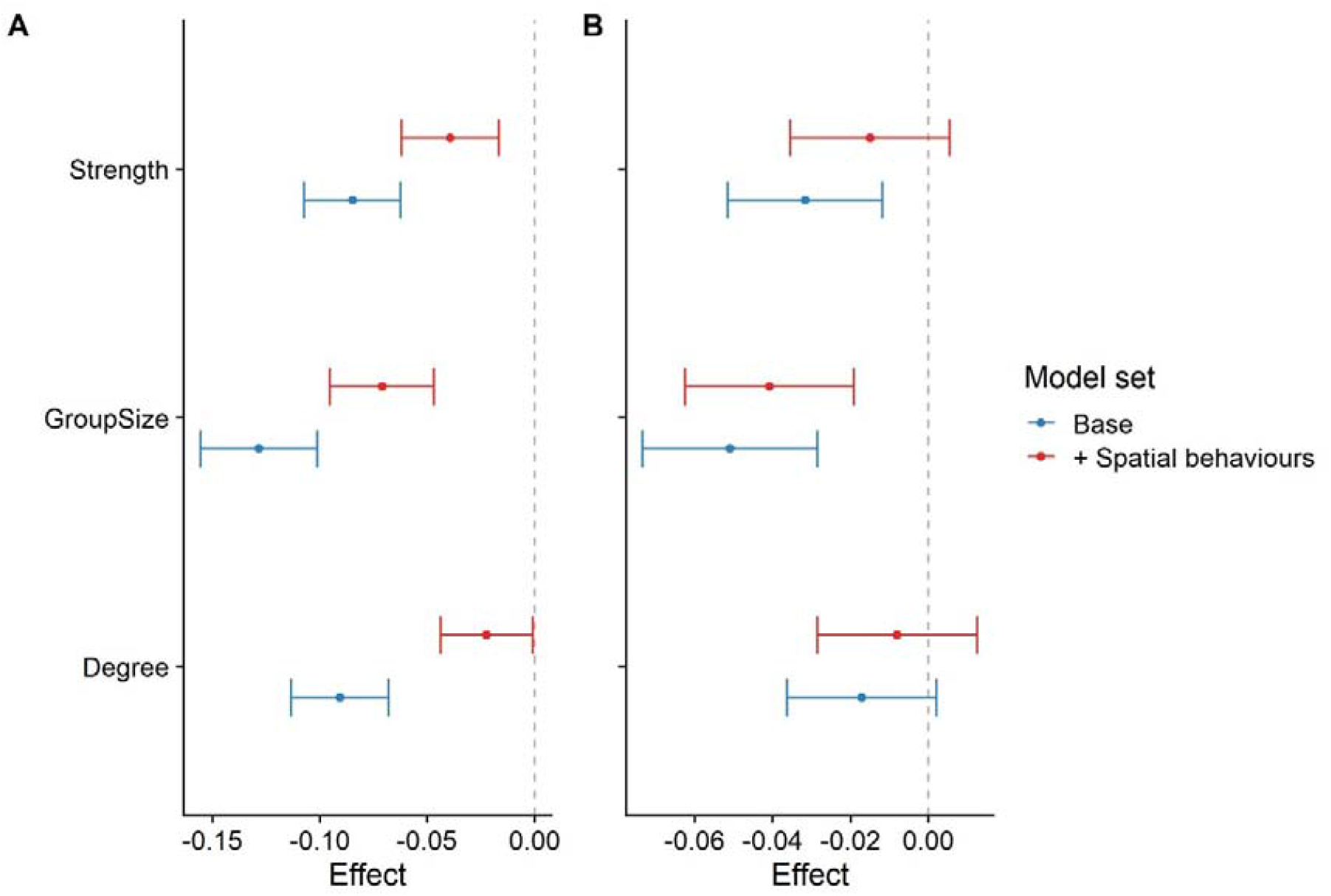
Model estimates for age effects on sociality in our longevity models (blue colours; Model Set 2); and in the same models with spatial behaviours accounted for (red colours; Model set 6). Dots represent the mean of the posterior effect estimate distribution; error bars denote the 95% credibility intervals of the effect. Panel A displays the effects without an SPDE effect fitted; panel B displays the same effects with SPDE fitted.

**Supplementary Figure 4:**
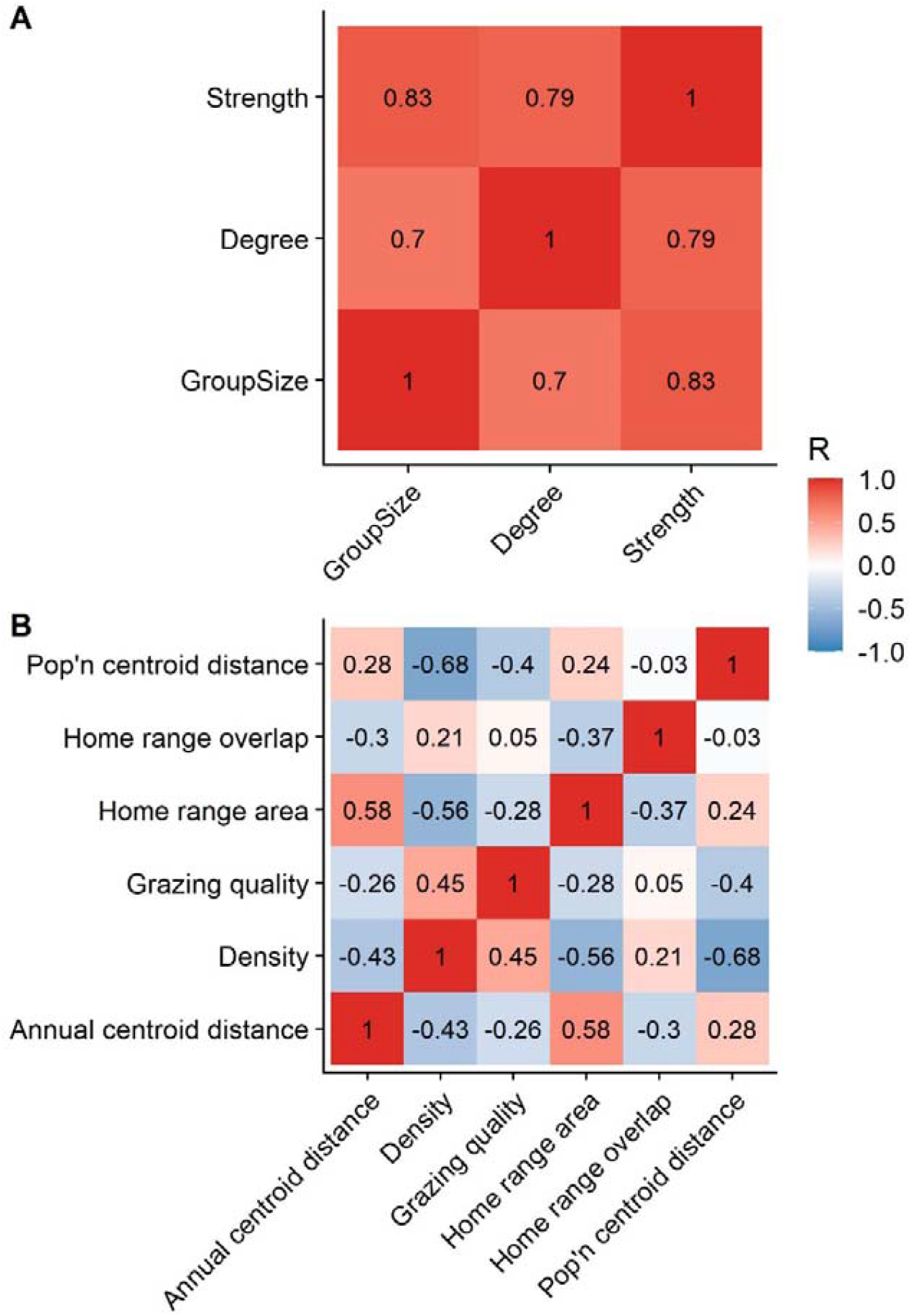
Pairwise correlations among social (A) and spatial (B) behavioural metrics. Panels that are more blue are more negative; panels that are more red are more positive.

